# Evidence of progenitor cell lineage rerouting in the adult mouse hippocampus

**DOI:** 10.1101/2020.06.08.140467

**Authors:** Daniela M.S. Moura, Juliana Alves Brandão, Celia Lentini, Christophe Heinrich, Claudio M. Queiroz, Marcos R. Costa

## Abstract

Cell lineage in the adult hippocampus comprises multipotent and neuron-committed progenitors. In the present work, we fate-mapped neuronal progenitors using Dcx-CreERT2 and CAG-CAT-EGFP double-transgenic mice (cDCX/EGFP). We show that three days after tamoxifen-mediated recombination in cDCX/EGFP adult mice, GFP+ cells in the dentate gyrus co-expresses DCX and about 6% of these cells are proliferative neuronal progenitors. After 30 days, 20% of GFP+ generated from these progenitors differentiate into GFAP+ astrocytes. Administration of the chemoconvulsants kainic acid (KA) or pilocarpine (PL) led to a significant increase in the number of GFP+ cells in both ipsi and contralateral dentate gyrus. However, while PL favored the differentiation of neurons in both ipsi- and contralateral sides, KA stimulated neurogenesis only in the contralateral side. In the ipsilateral side, KA injection led to an unexpected increase of astrogliogenesis in the Dcx-lineage. These different effects of KA and PL in the Dcx-lineage are associated with distinct alterations of the parvalbuminergic plexus and inflammatory responses in the hippocampi. Finally, we also observed a small number of GFP+/GFAP+ cells displaying radial-glia morphology ipsilaterally 3 days after KA administration, indicating that Dcx-progenitors could regress to a multipotent stage. Altogether, our data suggest that cell lineage in the adult hippocampus is not unidirectional and can be modulated by environmental signals.

## Introduction

Generation and functional integration of new neurons occurs at discrete sites of the adult central nervous system (for review, see Lledo et al., 2006). In the adult rodent hippocampus, new granule cells (GCs) are constantly added to the dentate gyrus (DG) throughout life (van Praag et al., 1999), though at significantly lower levels in aging brains (Encinas & Sierra, 2012). Adult hippocampal neurogenesis contributes to learning and memory (for review, see Aimone et al., 2014) due to the unique responses of immature GCs to activity patterns entering the DG (Marín-Burgin et al., 2012). It also provides a powerful model to study the integration of newly generated neurons into preexisting matured neuronal circuits (Sailor et al., 2017).

Generation of new GCs in the adult hippocampus follows a stereotyped cell-lineage progression, with multipotent radial-glia like cell (RGCs, Type 1) progenitors generating glia-like non-radial (Type 2a) cells, which in turn engender neuronal-committed progenitors (Type 2b and 3), directly responsible for the production of post-mitotic neurons (Steiner et al., 2006). RGCs can transit from a quiescent to an activated state by environmental signals (Dumitru et al., 2017; Song, M. Christian, et al., 2012) and retain the ability to differentiate into astrocytes (Bonaguidi et al., 2011; Encinas et al., 2011). However, it is largely believed that RGCs either follow an astroglial-lineage producing new RGCs and astrocytes or follow a neuronal-lineage, comprising Type 2b and 3 progenitors, which terminates with the generation of new granule neurons (Bonaguidi et al., 2011; Dumitru et al., 2017; Encinas et al., 2011; Steiner et al., 2004; Suh et al., 2007). Network activity affects cell proliferation and differentiation in the adult DGs and can be modulated by exposure to enriched environments (EE), running wheels, learning paradigms, stress and pathological conditions, such as epilepsy (Aimone et al., 2014). These modulations in cell lineage progression are likely mediated through different gamma-aminobutyric acid (GABA) levels in the DG acting on the gamma-2 subunit of the GABA type A receptor expressed in RGCs (Song, Zhong, et al., 2012). GABA activity promotes neuronal differentiation and stem cell quiescence, whereas reduced GABA activity favors RGCs self-renewal and differentiation to astrocytes (Dumitru et al., 2017; Song, Zhong, et al., 2012). Interestingly, the effect of GABA in RGCs is likely due to diffusion from nearby synapses (Song, Zhong, et al., 2012). However, activation of hilar interneurons elicits a GABA_A_ receptor-sensitive response in Type 2 progenitors, causing depolarization and opening of voltage-dependent calcium channels (Tozuka et al., 2005). Calcium influx promotes NeuroD expression and leads to neuronal differentiation (Deisseroth et al., 2004).

Intrahippocampal injection of kainic acid (KA) in rodents is used to model mesial temporal lobe epilepsy (MTLE) and is associated with a transient increase in cell proliferation, depletion of RGCs and astrogliogenesis in the DGs (Danzer, 2018 and references herein). Fate mapping of Nestin-expressing cells shows that KA-induced seizures lead to activation of RGCs and their conversion into reactive astrocytes, which exhausts adult hippocampal neurogenesis in the long-term (Sierra et al., 2015). However, Nestin is also expressed in Type 2 progenitors, and there is the possibility that KA could be inducing a lineage shift in these cells.

To address this possibility, we used a Dcx-CreERT2 transgenic mouse line to fate map the lineage of DCX-expressing cells. We show that DCX-expressing cells contribute a small proportion of astrocytes in the DG, even under physiological conditions. Increased network activity induced by KA or pilocarpine (another commonly used chemoconvulsant with different mechanism of action; (Moura et al., 2019)) has different effects on DCX-progenitors. While KA shifts the lineage towards an astrogliogenic fate within the injection side, pilocarpine enhances neurogenesis. Increased neuronal activity in the DG contralateral to the injection site of both pilocarpine and KA leads to augmented neurogenesis. Lastly, we demonstrate divergent effects of these drugs in the degeneration of DG parvalbumin-expressing neurons, further supporting the notion that GABAergic signaling controls the lineage progression of adult hippocampal neural progenitor cells.

## Results

### Dcx-lineage in the adult hippocampus encompasses astrocytes

Cell lineage in the adult hippocampus comprises multipotent (Type 1 and 2a) progenitors and neuron-determined (Type 2b and 3) progenitors (Steiner et al., 2004). To label and follow the latter, we generated double-transgenic mice crossing Dcx-CreERT2 (Zhang et al., 2010) and CAG-CAT-EGFP mice (Nakamura et al., 2006), hereafter referred to as cDcx/GFP. Animals and were killed 3, 7, or 30 days after tamoxifen (TAM) treatment (Figure 1A). Using confocal microscopy, we independently analyzed the co-localization between GFP and DCX, GFP and GFAP or GFP and NEUN (Figure 1B-J). Three days after TAM administration, 97% of GFP+ cells in the dentate gyrus (DG) co-expressed the protein DCX and were either cells with short horizontal processes located in the subgranular zone (SGZ) or cells with radially oriented processes located in the inner half of the granular layer (Figure 1B-D, K; n=5 animals; 106 GFP+/DCX+ out of 108 GFP+ cells), as expected for type 2 progenitors and immature neurons, respectively (Brown et al., 2003; Jagasia et al., 2009). In contrast, only 0.5% of GFP+ cells co-stained for GFAP in the hilus (Figure 1 K; n=5 animals; 1 GFP+/GFAP+ cell out of 198 GFP+ cells). Seven days after recombination, 90% of GFP+ cells occupied the inner half of the granular layer, showed morphologies reminiscent of immature granule neurons and co-expressed DCX (Figure 1K; n=4 animals; 368 GFP+/DCX+ out of 415 GFP+ cells). About 1-2% of GFP+ cells already expressed NEUN (Figure 1K), a protein associated with the maturation of adult-born neurons in the DG. Interestingly, we also observed at this time-point a small but consistent population of GFP+ cells expressing the astrocytic protein GFAP (Figure 1E-G, K; n=4 animals; 13 GFP+/GFAP+ out of 476 GFP+ cells). Thirty days after recombination, 15% of GFP+ cells still expressed DCX and 82% of the recombined cells displayed typical morphology of mature granule neurons and expressed NEUN or CTIP2 (Figure 1K; n=4 animals; 26 GFP+/DCX+ and 142 GFP+/CTIP2 or NeuN+ out of 172 GFP+ cells). At this time-point, we could still detect about 3% of GFP+ cells expressing GFAP (Figure 1K; n=4 animals; 4 GFP+/GFAP+ out of 131 GFP+ cells). Altogether, these data confirm previous observations indicating that the Dcx-lineage in the adult hippocampus encompasses mostly newly generated granule neurons (Zhang et al., 2010), but unexpectedly contains a small percentage of astrocytes at days 7 and 30.

**Figure 1.**
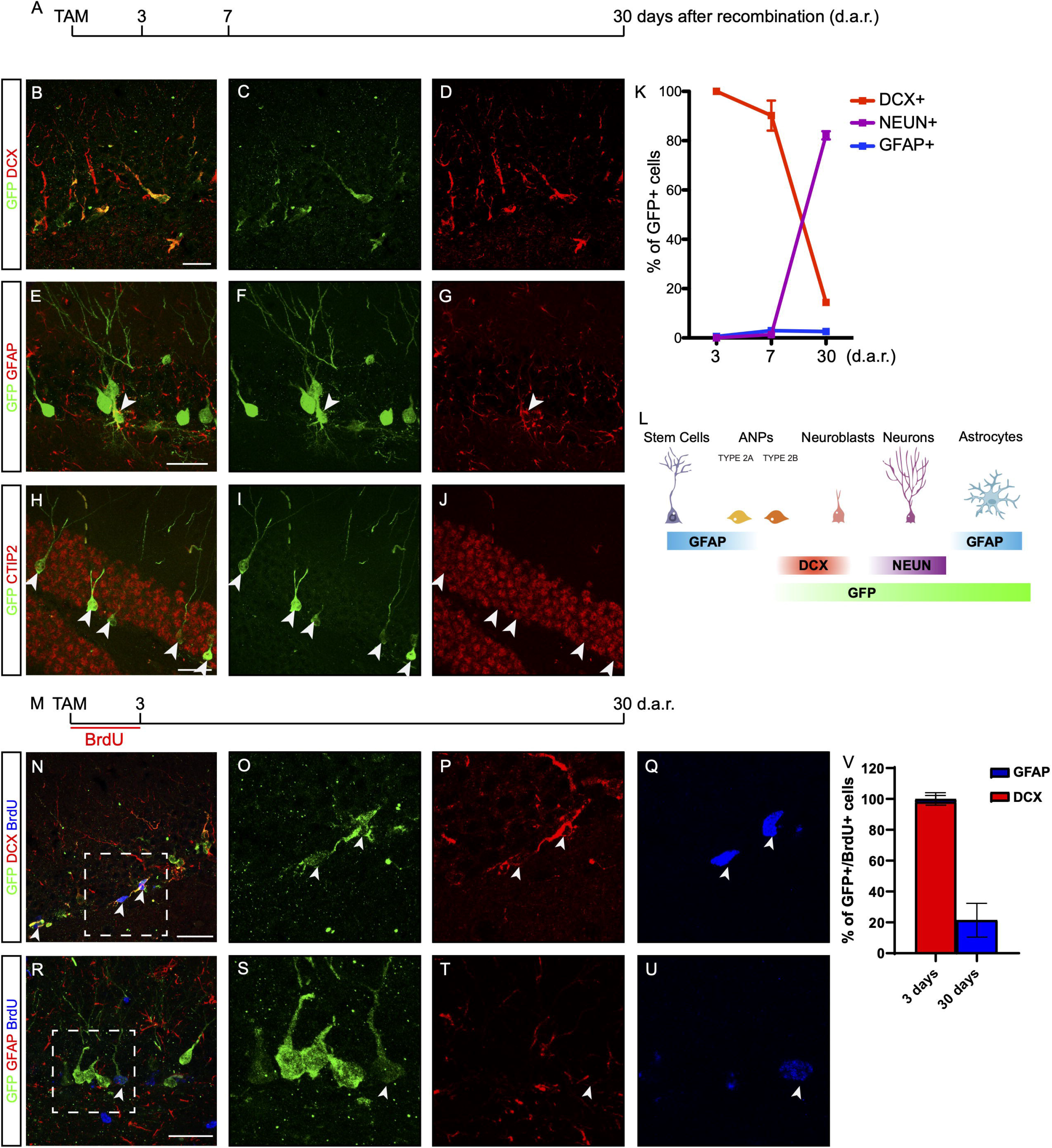
Dcx-lineage comprised mostly neurons and a small fraction of astrocytes. (A) Timeline presenting timepoints in which animals were killed for observation and quantification of labeled cells after TAM injection. (B-J) Coronal sections of the hippocampus of cDCX/EGFP animals immunolabeled for GFP (green), DCX, GFAP or Ctip2 (red) after tamoxifen treatment. Observe that GFP expression is restricted to small DCX+ cells in the SGZ 3 days after tamoxifen injection (B-D). (E-G) A second population of GFP+/GFAP+ cells showing morphologies of astrocytes is noticed 30 days after tamoxifen administration (arrowheads). (H-J) After 30 days cells present mature granule cell morphology, express neuronal marker NEUN. (K) Graphic showing the quantification of GFP cells expressing makers of different cell populations (n=4 to 5 animals per timepoint). (L) Schematic representation of hippocampal neurogenesis showing main markers in each phase of cell development. (M) Timeline presenting experimental protocol from group receiving TAM injection followed by 3 days of BrdU administration on water. Animals were perfused 3 or 30 days after recombination (d.a.r.). (N-Q) Expression of DCX in GFP+/BrdU+ cells 3 d.a.r. (R-U) 30 days after Dcx-mediated recombination, GFP+/BrdU+ are rare possibly due to BrdU dispersion. (V) Graphic showing the quantification of GFP+/BrdU+ cells expressing DCX or GFAP. (n=5 animals in “3 days” group; n=3 in “30 days” group). ML - Molecular Layer; GCL - Granular Cell Layer. Scale bar: 20um.

To directly probe whether GFP+ astrocytes could be derived from Dcx-expressing progenitors in the DG, we treated cDcx/GFP adult animals with tamoxifen followed by 3-days BrdU treatment (Figure 1M). A subset of animals was killed immediately after BrdU administration (therefore, 3 days after tamoxifen), and the remaining animals were allowed to survive for an additional 27 days (30 days after tamoxifen). We found that 94% of recombined GFP+ cells did not incorporate BrdU during the 3-days treatment (503 GFP+/BrdU-out of 534 GFP+ cells). In contrast, the remaining 6% of GFP+ cells were labeled with BrdU, indicating that only a small fraction of recombined cells in the DG of cDcx/GFP mice are progenitors. Among those cells, 94% were also DCX+ (Figure 1V; n=5 animals; 29 GFP+/BrdU+/DCX+ out of 31 GFP+/BrdU+ cells). In contrast, only in 1 out of 5 animals analyzed we could detect a single GFP+/BrdU+ cell co-expressing GFAP (Figure 1V; n=5 animals; 1 GFP+/BrdU+/GFAP+ out of 51 GFP+/BrdU+ cells), indicating that unspecific recombination GFAP-expressing progenitors in the DG of cDcx/GFP mice following our TAM administration protocol is unlikely. Even considering the total population of BrdU+ cells in the DG that did not express GFP, only 4% of cells coexpressed GFAP, which is in accordance with the low number of Type 1/2a progenitors in the DG (Encinas et al., 2011). Notably, however, 30 days after Dcx-mediated recombination, we observed that up to 19% of GFP+/BrdU+ cells co-expressed the astrocyte protein GFAP (Figure 1; n=3 animals; 5 GFP+/BrdU+/GFAP+ out of 27 GFP+/BrdU+ cells). Altogether, these observations may suggest that at least some of GFP+/BrdU+/DCX+ cells, 3 days after recombination, are DCX+ Type 2b/3 neuronal progenitors that retain the ability to generate astrocytes.

### Divergent effects of kainic acid and pilocarpine on the Dcx-lineage

The observation that a small subset of neuronal progenitors in the Dcx-lineage could retain the potential to generate astrocytes prompt us to evaluate whether changes in network excitability could interfere with the lineage progression of these progenitors, as it has been shown for other systems (Lin et al., 2017). To that, we employed chemoconvulsants traditionally used to trigger dentate gyrus network reorganization in studies modeling the human mesial temporal lobe epilepsy (MTLE) in rodents (Lévesque et al., 2016). It has been previously shown that the administration of chemoconvulsants increases astrogliogenesis from multipotent Nestin-expressing progenitors (Sierra et al., 2015) and neurogenesis from Dcx-expressing cells (Moura et al., 2019). Unilateral intrahippocampal injection of kainic acid (KA, 50 nL, 20 mM; Cayman Chemical) in 2 months-old cDcx/GFP animals induced long-lasting bilateral epileptiform discharges (Figure S1) and behavioral seizures indicative of *status epilepticus* (SE), as previously described (Moura et al., 2019). According to previous data in the literature suggesting that cell proliferation/survival is affected by chemoconvulsants (Kralic et al., 2005; Heinrich et al., 2006; Moura et al., 2019), we observed an overall increase in the number of GFP+ cells in both hippocampi of KA- and PL-treated animals (Figure S2), and severe granule cell dispersion restricted to the KA-ipsilateral side (Figure S1). To fate-map the Dcx-lineage in KA-injected cDcx/GFP animals, tamoxifen was administered one week after SE, and the phenotype of GFP+ cells was evaluated 4 weeks later (Figure 2). In the contralateral side of the KA injection, despite the increased electrical activity detected after KA injection (Figure S1), we observed that 94 ± 2% of GFP+ cells displayed typical granule cell morphology and expressed CTIP2, a proportion slightly higher than the 86 ± 10% observed in saline-injected control animals (Figures 2B, D-E, H; Figure S3; n vehicle=3 animals; 243 neurons out of 285 GFP+ cells in contralateral; n KA=5 animals; 1410 cells out of 1510 GFP+ cells in contralateral side). Surprisingly, however, we found that 90 ± 7.5% of GFP+ cells in the ipsilateral KA injection side displayed astrocytic morphologies and expressed GFAP, with the remaining 10% of GFP+ cells differentiating into neurons (Figures 2F-G, I; N vehicle = 3 animals, 315 neurons out of 369 GFP+ cells; n KA = 5 animals; 113 neurons out of 1210 GFP+ cells). In sharp contrast, the proportion of GFP+ cells co-expressing GFAP was about 2% in both ipsi and contralateral hippocampus of animals treated pilocarpine (PL), whereas the proportion of neurons increased up to 98% (Figure 2C, H-I; n PL=3 animals; 1616 neurons out of 1658 GFP+ cells in contralateral and 2317 neurons out of 2364 GFP+ cells in ipsilateral side). Similar to KA, intrahippocampal injection of 700nl of PL (700 μg/μL; Sigma-Aldrich) induces bilateral epileptiform electrical activity and behavioral changes indicative of *status epilepticus* (SE) (Moura et al., 2019). Together with our finding of increased neurogenesis in the contralateral side of KA-injected animals, these observations in PL-treated animals suggest that increased electrical activity could be permissive to neurogenesis, whereas the increased astrogliogenesis observed in the ipsilateral hippocampus of KA-treated animals (Figure 2; Sierra et al., 2015) may rely on other local effects of this chemoconvulsant.

**Figure 2.**
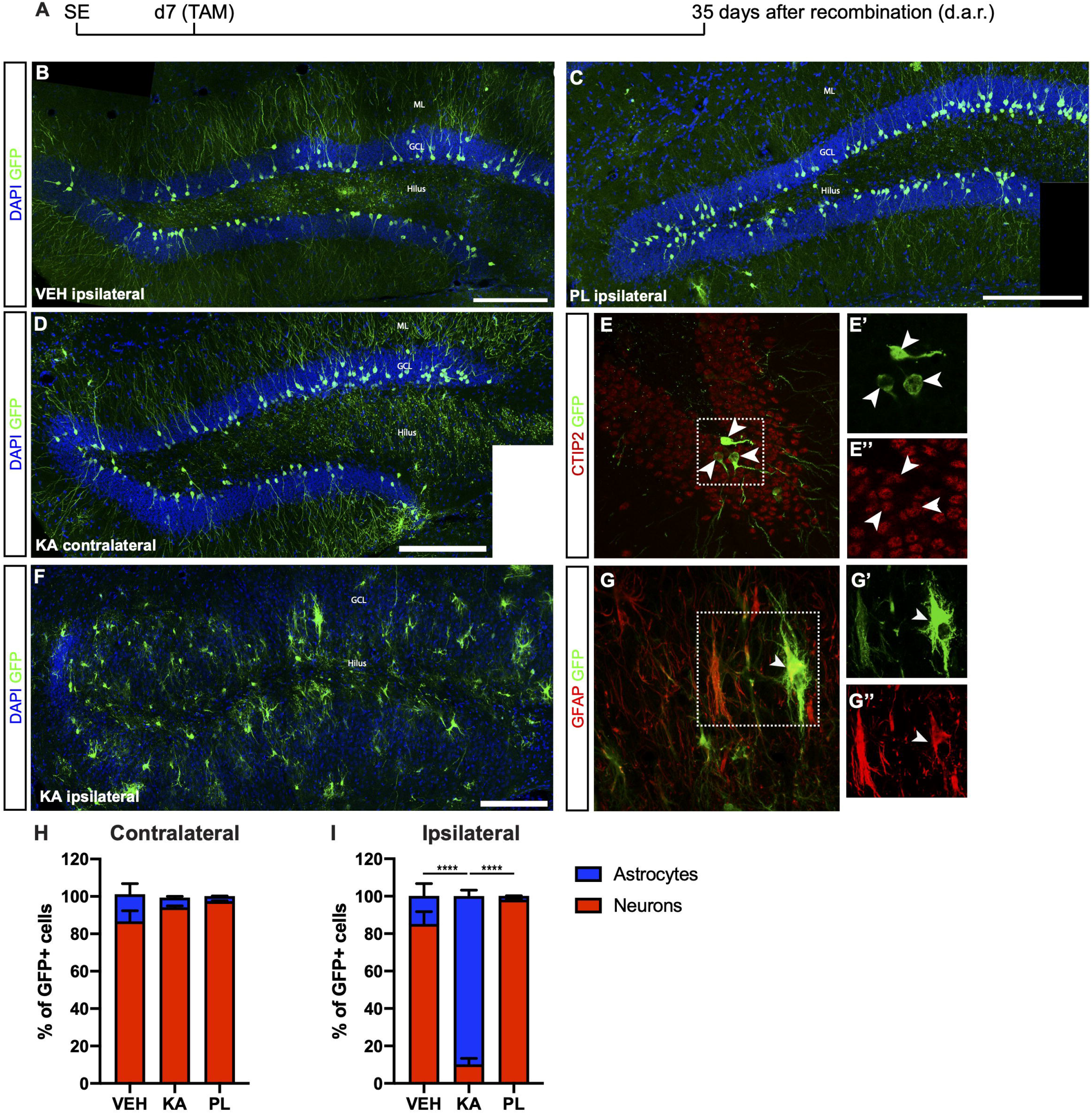
Kainic acid and pilocarpine have opposite effects in Dcx-lineage progression. (A) Timeline presenting experimental protocol from group receiving two TAM injections 7 days after SE. Animals were perfused 35 days later. (B-D) Mosaic composition from vehicle (B) and PL (C) ipsilateral side and KA contralateral hippocampus (D) showing GFP+ granule cells with typical morphology. (E) Confocal image of contralateral hippocampus in KA group showing the expression of CTIP2 in GFP+ granule cells in single plane z-projection images. (F) Ipsilateral hippocampus of KA injected animals displays severe GCD and most GFP+ cells have morphologies of reactive astrocytes. (G) Confocal image of ipsilateral DG showing the expression of GFAP in GFP+ cells in single plane z-projection images. (H-I) Quantification of GFP+ cells differentiating into granule neurons (H) or astrocytes (I) in the contralateral and ipsilateral sides of KA-injected animals. ML - Molecular Layer; GCL - Granular Cell Layer. (n vehicle=3; n KA=5; n PL=3; two-way ANOVA followed by Tukey’s multiple comparisons: ****Adjusted p value (KA vs. VEH and KA vs. PL) < 0.0001). Each value represents the mean ± SEM. Scale bars: 100um (B-D, F) and 20um (E,G).

To rule out the possibility that KA injection could induce activation of the Dcx promoter directly in astrocytes leading to CRE-ERT expression and subsequent TAM-mediated recombination, we injected KA in the striatum, close to the rostral migratory stream (RMS) of cDcx/GFP adult mice, followed by a single TAM injection 7 days later, and analyzed the expression of GFP in reactive astrocytes after 35 days. In contrast to the hippocampus, no GFP+/GFAP+ cells were observed in the striatum or olfactory bulb (OB; Figure S4), supporting the notion that KA *per se* does not activate the Dcx promoter in astrocytes of cDcx/GFP double-transgenic animals. Also consistent with the interpretation that Dcx-promoter is active only in the neuronal lineage of the RMS-OB system, we observed that GFP+ cells differentiated into OB granular and peri-glomerular interneurons 35 days after TAM (Figure S4).

### KA induces astrogliogenesis from Dcx-progenitors recombined 1 day before SE

Next, to unambiguously exclude possible influences of KA on the activity of the Dcx promoter in the hippocampus, we recombined Dcx-expressing cells before SE induction. To that, cDcx/GFP mice received tamoxifen 1 day before intrahippocampal injection of KA or vehicle (Figure 3). Animals also received BrdU for three consecutive days to label proliferating cells and were killed immediately after this period or 30 days after KA. We observed that the proportion of GFP+ cells undergoing cell proliferation during the 3 days BrdU-treatment was similar between control and KA animals (Figure 3B; n vehicle=5 animals, 268 GFP+ cells in the contralateral and 281 in the ipsilateral side; n KA=3 animals, 249 GFP+ cells in the contralateral and 759 in the ipsilateral side). However, while in both hippocampi of control animals and in the contralateral side of KA-treated animals, virtually all GFP+/BrdU+ cells co-expressed DCX, about 14% of GFP+/BrdU+ cells already expressed GFAP in the ipsilateral side of KA-injected mice at this time point (Figure 3C; n vehicle=5 animals, 21 GFP+/BrdU+ cells in the contralateral and 18 in the ipsilateral side; n KA=3 animals, 14 GFP+/BrdU+ cells in the contralateral and 48 in the ipsilateral side). This fraction of GFP/GFAP/BrdU cells in the KA-ipsilateral side further increased to about 58% after 30 days (Figure 3D; n vehicle=4 animals, 18 GFP+/BrdU+ cells in the contralateral and 12 in the ipsilateral side; n KA=3 animals, 25 GFP+/BrdU+ cells in the contralateral and 46 in the ipsilateral side). Also, despite the complete absence of GFP+/GFAP+/BrdU+ at day 3 in control animals and in the KA-contralateral side, these cells could be observed at small but consistent frequencies 30 days after recombination (Fig 3D-E). Altogether, these observations confirm that astrocytes are encompassed within the lineage of Dcx-expressing Type 2b/3 cells and that generation of new astrocytes from Type 2b/3 cells depends on cell proliferation.

**Figure 3.**
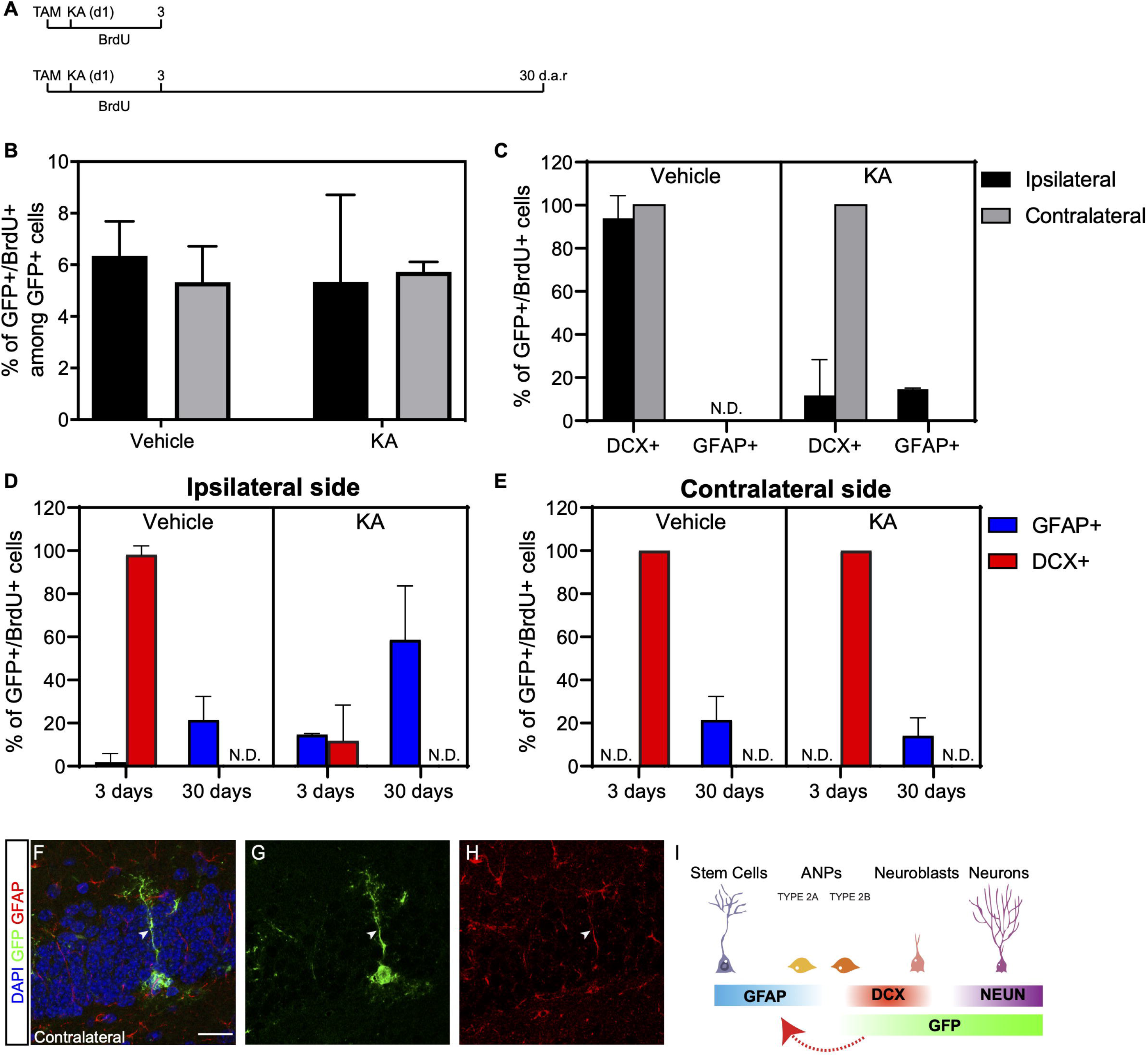
DCX-cells retain the capacity to proliferate and generate astrocytes after 1 month. (A) Timeline representing experimental protocols in which one tamoxifen injection was administered 1 day before SE induction with KA. Animals received BrdU in water during 3-days and were perfused right after this period, or 30 days after recombination. (B-C) Quantification of proliferative GFP+ cells after 3 days (B), and expression of DCX or GFAP among these cells (C) on ipsilateral and contralateral side of vehicle and KA injected animals. (D-E) Quantification of proliferative GFP+ cells after 3 and 30 days in ipsilateral side (D) and contralateral (E) expressing DCX or GFAP. (F-H) Example of GFP+/GFAP+ cell (RGL) in the SGZ of KA injected cDCX/EGFP animals 3 days after recombination. (I) (A) Schematic representation of the evidence supporting a possible regression (dotted red arrow) of recombined Type 2B/ DCX-expressing progenitors to a Type 1/ RGL/ GFAP-expressing multipotent progenitor state in the adult hippocampus.

The generation of astrocytes from Dcx-expressing progenitors (Type 2b and 3) may suggest that those cells are bipotent or may reverse to a more primitive stage in the lineage when cells retain astrogliogenic potential (Bonaguidi et al., 2011; Encinas et al., 2011). To directly address this second possibility, we set out to investigate whether cells within the Dcx-lineage could adopt hallmarks of Type 1, radial glia-like (RGL) progenitors. In both untreated, vehicle- and PL-injected animals we were unable to detect any GFP+/GFAP+ RGL cells in the SGZ at all time points analyzed (3, 7 or 30 d.a.r). In contrast, we found a small number of GFP+/GFAP+ RGL cells in animals treated with KA both in the ipsilateral and contralateral hippocampus (Figure 3G; n=3 animals; 8 GFP+/GFAP+ RGL cells out of 387 GFP+ cells). These observations could suggest that Type 2b Dcx-expressing progenitors are not fully committed to differentiate into neurons but may rather return to more primitive stages in the lineage and resume multipotency under non-physiological conditions.

### Progenitors in the Dcx-lineage retain the ability to generate neurons and astrocytes 1 month after recombination

The interesting observation that Dcx-expressing progenitors can regress to previous stages of the lineage and generate astrocytes prompted the question as to which extent this potential could be observed in progenitors recombined at earlier time-points before KA injection. To address this question, we injected tamoxifen in cDcx/GFP adult mice 35 days before intrahippocampal injections of chemoconvulsants (Figure 4). Fifteen days after KA treatment, we observed a significant increase in the proportion of astrocytes among GFP+ cells in the KA-ipsilateral side as compared to the contralateral-KA side, PL- and vehicle-injected animals (Figure 4B-C, F; n vehicle=3 animals, 65 astrocytes out of 974 GFP+ cells in contralateral and 57 out 988 GFP+ cells in ipsilateral side; n PL=3 animals, 34 astrocytes out of 474 in contralateral and 38 out of 363 GFP+ cells in ipsilateral side; n KA=4 animals; 160 astrocytes out of 1053 in contralateral and 405 out of 1170 GFP+ cells in ipsilateral side). Yet, and consistent with our previous findings, a small population of GFP+/GFAP+ cells could be observed in both hippocampi of PL and control animals, as well as in the contralateral-KA side (Figure 4F). These observations suggest that Dcx-expressing progenitors recombined 5 weeks before KA injection retain their ability to proliferate and generate astrocytes. In fact, when the same experiment was performed with the additional treatment of animals with BrdU for the whole period after KA-injection, we observed that most GFP+ astrocytes in the KA-ipsilateral side incorporated BrdU (Figure 4G; n KA = 3 animals; 55 GFP+/GFAP+/BrdU+ out of 77 GFP+/GFAP+ cells), indicating that despite the low numbers of progenitor cells recombined within the Dcx-lineage, some are still active 35 days after recombination and retain the potential to generate astrocytes. According to this notion, we also detected GFP+/GFAP+ RGL cells in the SGZ of these animals (Figure S5).

**Figure 4.**
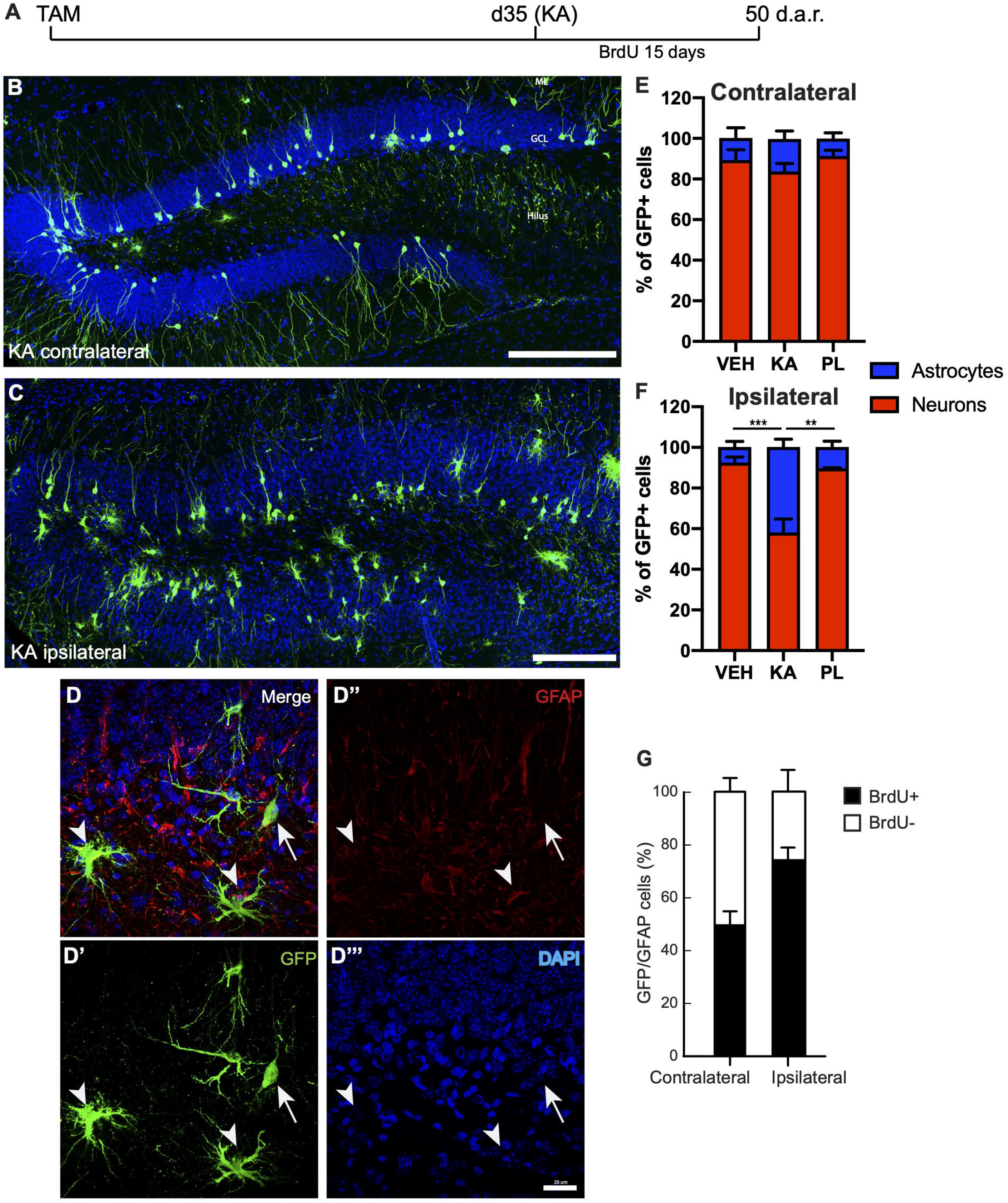
Kainic acid injection enhances astrogliogenesis from previously recombined DCX-cells. (A) Timeline representing experimental protocol where two tamoxifen injections were administered 35 days before SE induction with KA. Animals received BrdU during the whole period after KA and were perfused two weeks later. (B) Coronal confocal mosaic image of the contralateral dentate gyrus shows the majority of GFP+ cells with typical neuronal morphologies. (C) The image of ipsilateral dentate gyrus shows GCD and many GFP+ cells with morphologies of astrocytes. (D) Examples of GFP+/GFAP+ astrocytes (arrowheads) and neurons (arrows) in the ipsilateral DG. (E-F) Quantification of GFP+ cells differentiating into neurons (E) and astrocytes (F) in the ipsi and contralateral sides of controls and KA-treated animals. (G) Quantification of proliferative GFP+/GFAP+ cells in both hemispheres in KA injected animals ML - Molecular Layer; GCL - Granular Cell Layer. Scale bars: 50um (B-C) and 20um (D). (n vehicle=4; n KA=4; n PL=3; two-way ANOVA followed by Tukey’s multiple comparisons: **Adjusted p value (KA vs. PL) = 0.0018; ***Adjusted p value (KA vs. VEH) = 0.0008).

Next, we followed an independent approach based on retroviral-mediated labeling of fast-dividing types 2/3 intermediate progenitors in the hippocampal subgranular zone (Figure S6). To this end, adult mice were initially injected with a retrovirus encoding a DsRed fluorescent reporter one month before receiving intrahippocampal injection of KA or saline for controls, and the identity of DsRed+ cells was analyzed a month later (Figure S6A).

Interestingly, we could also observe that injection of KA, 4 weeks after retroviral infection, shifted the lineage towards a gliogenic fate (Figure S6B-D; KA: 567 glia out of 1045 DsRed+ cells in the ipsilateral side, n = 7 mice; vehicle: 21 astrocytes out of 224 DsRed+ cells in the ipsilateral side, n = 4 mice). Altogether, these data using two independent strategies to fate-map the progeny of type2b/3 progenitors further support our interpretation that these cells retain an astrogliogenic potential.

### Loss of parvalbumin-expressing interneurons correlates with increased astrogliogenesis

To get some insights about the possible mechanisms controlling the switch from neurogenesis to gliogenesis in the Dcx-lineage observed upon KA treatment, we set out to evaluate the effects of this drug and PL on the parvalbuminergic plexus of the DG. GABA activity mediated by parvalbumin (PV)+ interneurons promotes stem cell quiescence and neuronal differentiation in the adult hippocampus (Dumitru et al., 2017; Song, Zhong, et al., 2012). We hypothesized that KA and PL could differently affect PV+ interneurons, which could help to explain the divergent effects on cell lineage progression in the DG of mice treated with glutamatergic and cholinergic agonists, respectively. To test this possibility, we analyzed the hippocampus of adult mice 3 days after saline, KA or PL injection (Figure 5). According to previous data in the literature (Song, Zhong, et al., 2012), we observed a complex parvalbuminergic plexus within the subgranular zone, originating from PV+ cells in the hilus and in close relation with Type 2b BrdU+/DCX+ progenitors (Figure 5A). Next, we quantified the total number of PV+ interneurons and found that the ipsi/contralateral ratio of PV+ cells in the DG was significantly reduced in KA-injected animals, but not in animals treated with PL (Figure 5B; n=3 animals per group, 4-5 coronal sections from each animal). Interestingly, whereas the total number of PV+ cells in the ipsilateral DG of control and PL animals were similar, these cells were virtually absent in KA-injected animals (Figure 5C), in agreement with previous studies (Bouilleret et al., 2000). Conversely, the number of PV+ cells in the contralateral KA-side and in both hippocampi of PL-injected animals was similar to controls (Figure 5C). A similar pattern was observed when we counted all PV-expressing cells in the whole hippocampus, including CA region (Figure S7), suggesting that the loss of PV+ interneurons is restricted to the ipsilateral hippocampus of KA-treated animals. Thus, the region where Dcx-progenitors generate astrocytes (ipsilateral KA-injected hippocampus) shows both an increase of electrical activity (Figure S1) and overall reduction in the number of PV+ interneurons in the dentate gyrus.

**Figure 5.**
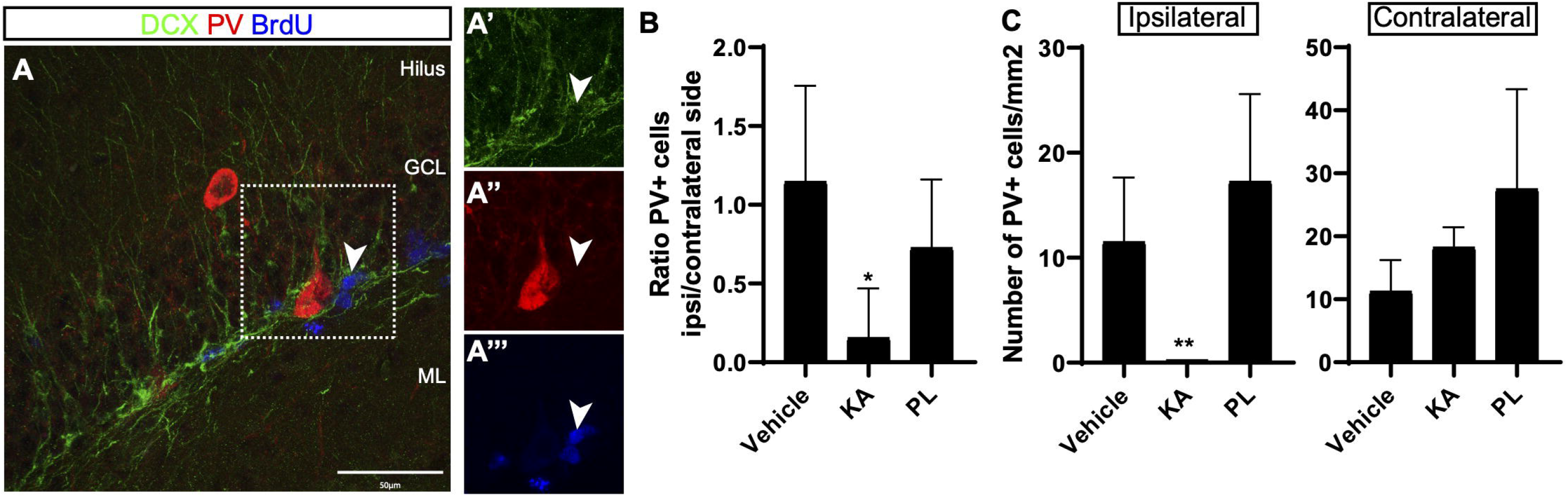
Parvalbuminergic plexus degenerate in the ipsilateral side of KA injection. (A) Coronal section of the hippocampus 3 days post injection (dpi) of vehicle. Note the presence of PV+ interneurons in the granule cell layer (GCL). Observe process of PV+ cell in close contact with proliferating BrdU+/DCX+ cell (arrowhead) in the subgranular zone (SGZ). Scale bar: 50 μm. (B) Ratio of PV+ cells in the ipsi and contralateral sides of the hippocampus in different treatment groups. (C) Quantification of the number of PV+ interneurons per mm^2^ of the dentate gyrus 3 days after injection of saline, KA or PL. (n vehicle=4; n KA=4; n PL=4; Statistics: (B) ANOVA F_(2,9)_= 4.529; p=0.0436; Tukey’s multiple comparisons test DF = 9; Adjusted p value (control vs KA) = 0.0363; (C) ANOVA F_(2,9)_= 8.347; p=0.0089; Tukey’s multiple comparisons test DF = 9; Adjusted p value (KA vs PL) = 0.0077)

### Microglia activation is associated both with increased hippocampal neurogenesis and astrogliogenesis

Exacerbated inflammatory response has been described in rodent models of mesial temporal lobe epilepsy (Heinrich et al., 2006; Moura et al., 2019; Sierra et al., 2015), which negatively affects neurogenesis (Monje et al., 2003; Sierra et al., 2015). To evaluate whether unilateral KA- and PL-intrahippocampal injections could elicit different degrees of inflammation in the ipsi and contralateral sides we quantified the total number of resting and activated microglial cells 3 days after drug injection in the whole DG, including the hilar region, granule cell and molecular layers (Figure 6). Activated microglia cells (aMGCs) were classified based on the morphology and size of the cell body (Davis et al., 2017). While most MGCs in control animals were resting, we observed a significant proportion of aMGCs both in the ipsilateral (86% aMGCs) and contralateral hippocampus (61% aMGCs) of KA-treated animals (Figure 6N-O; n=3 animals per group; total number IBA1+ cells in ipsi and contralateral sides, respectively: Vehicle – 367 and 336; KA – 241 and 186; PL – 103 and 77). In contrast, only a small fraction of microglial cells was activated after PL injection (20% and 11% in the ipsi and contralateral hippocampus, respectively), which could indicate the presence of a subtler inflammatory response compared to KA-treated animals. Thus, the aMGCs observed in the DG are either associated with increased neurogenesis (PL and contralateral KA groups) or with induced astrogliogenesis (ipsilateral KA group).

**Figure 6.**
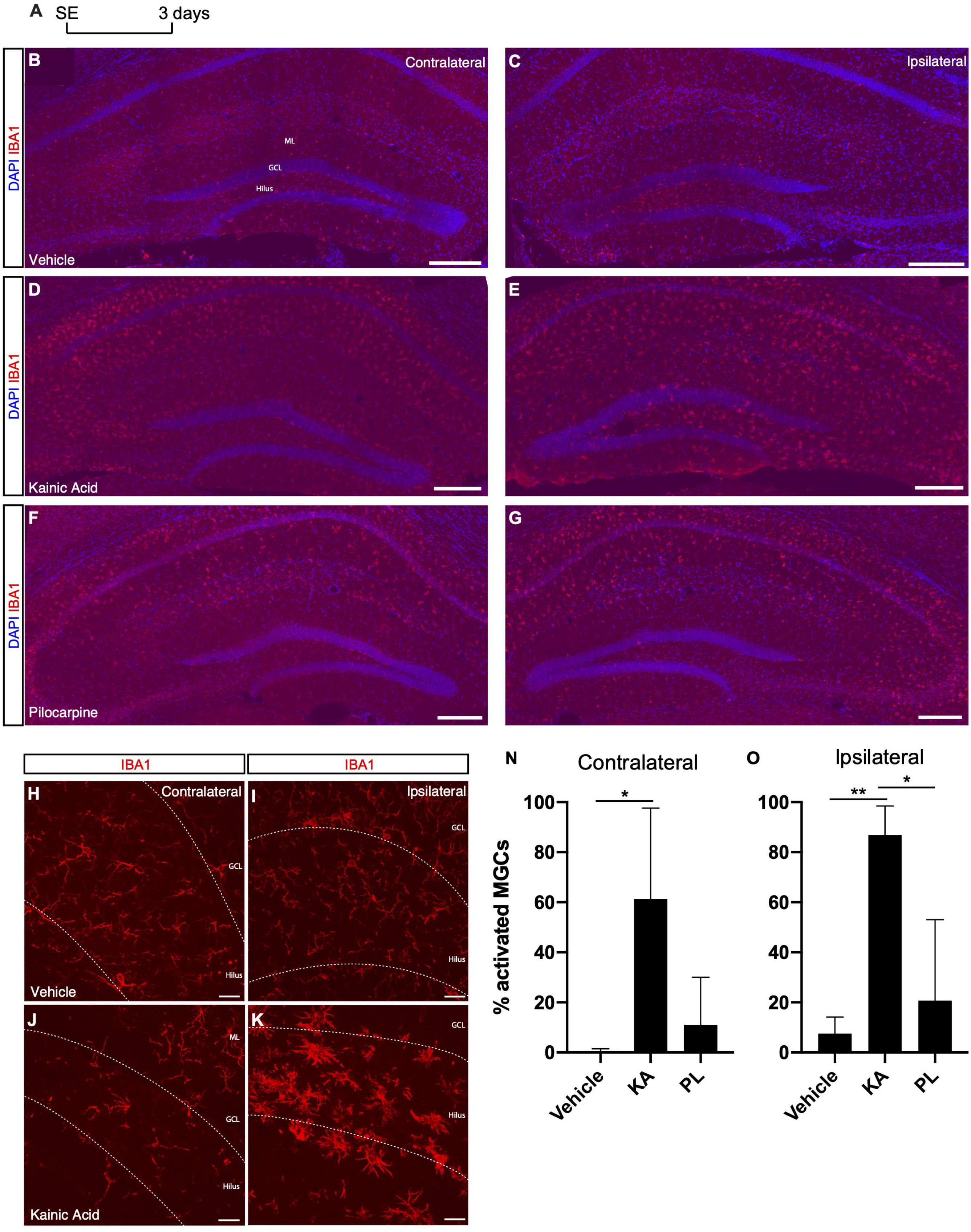
Microglia activation is stronger in the ipsilateral side of KA injection. (A) Timeline of experimental procedure to quantify Iba1+ cells. (B-C) Microglial cells in control group labeled with anti-Iba1 antibody and showing resting morphologies. (D-E) Increased signal for anti-Iba1 antibody in the KA group, indicating bilateral activation of microglia in the CA1/CA3 regions and in the ipsilateral DG hilus and molecular layer. (F-G) Increased signal for anti-Iba1 antibody in the PL group, indicating bilateral activation of microglia in the CA1/CA3 regions. (H-K) Higher magnification of the DG in vehicle (H-I) and KA (J-K) animals showing the morphological changes of microglial cells in the ipsilateral KA group. (N-O) Quantification of activated microglia in the ipsi or contralateral hippocampi of vehicle-, KA- and PL-treated animals. (n control = 3; n KA= 2; n PL= 3; Statistics: (N) ANOVA F_(2,5)_= 5.819; p=0.0495; Tukey’s multiple comparisons test DF = 5; Adjusted p value (control vs KA) = 0.0486; (O) ANOVA F_(2,6)_= 13.24; p=0.0063; Tukey’s multiple comparisons test DF = 6; Adjusted p value (vehicle vs KA) = 0.0072; (PL vs KA) = 0.0166).

## Discussion

Comprehensive knowledge about the intermediate steps enacted in the transition from an adult neural stem cell to neuronal state is critical to understand how neurons can be generated in the adult central nervous system. Here, we provide evidence that neurogenic cell lineage progression in the adult hippocampus is not a one-way process towards granule cell differentiation. Rather, intermediate progenitors retain the potential to regress to more primitive stages and regain astrogliogenic potential. Increased electrical activity induced by KA injection affects cell lineage progression in opposite directions depending on the concurring GABAergic activity and inflammatory status. While increased electrical activity in the ipsilateral DG, where degeneration of GABAergic PV+ interneurons and increased microglial activation leads to an increase in astrogliogenesis, in the contralateral side, where hilar GABAergic PV+ interneurons are preserved, augmented electrical activity further enhances neurogenesis. The latter effects are also correlated in the PL-injected animals, further supporting the notion that intermediate progenitors are responsive to neuronal network activity to controlling adult neurogenesis in the hippocampus (Deisseroth et al., 2004). We suggest that intermediate progenitors may function as sensors of global network activity and, therefore, serve as checkpoints for population lineage progression in the adult DG (Figure 7).

**Figure 7.**
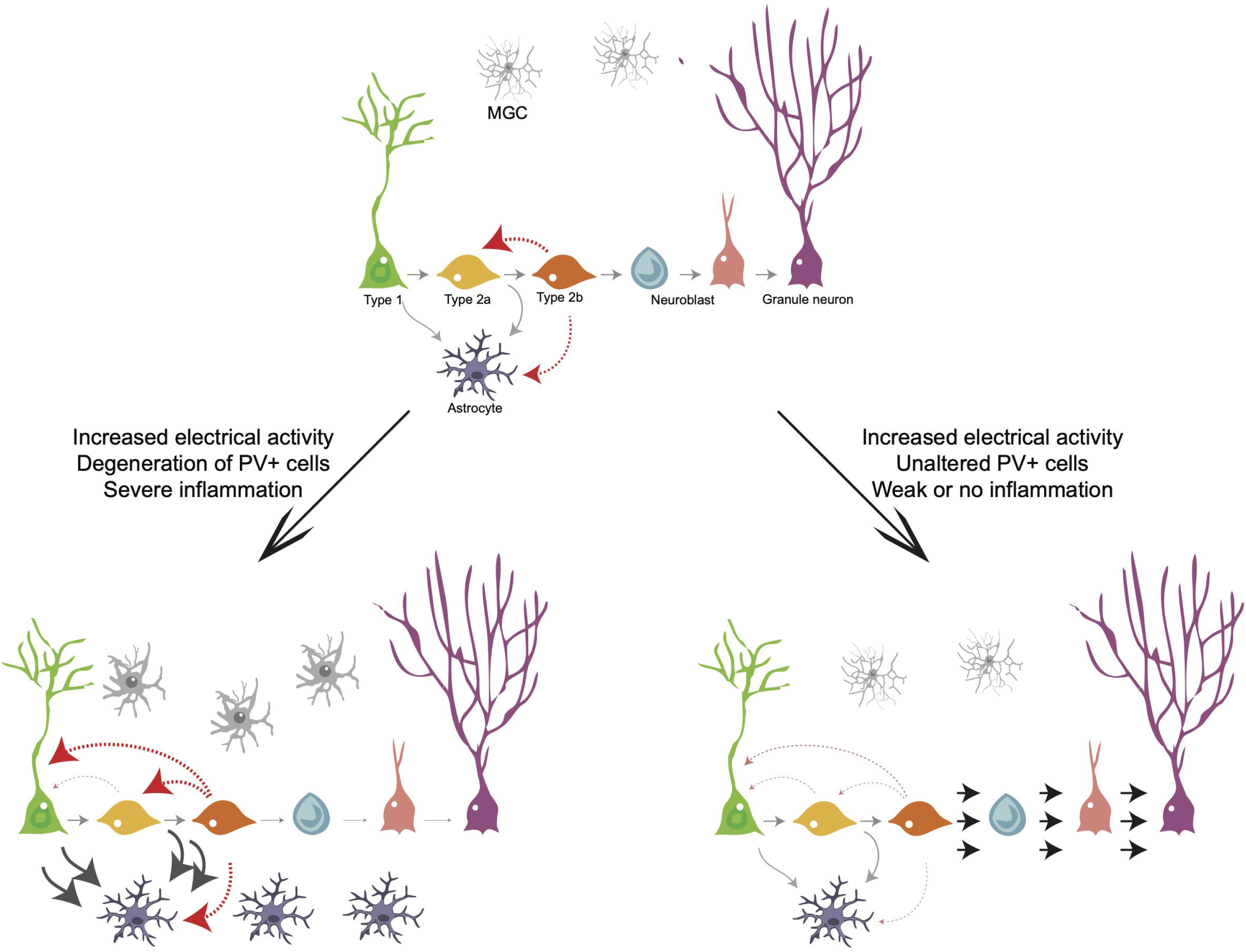
Cell lineage progression model in the adult hippocampus. (A) Grey arrows indicate previously described transitions during cell lineage progression from RGC to neurons and from RGC to astrocytes in the adult hippocampus. Red arrows indicate the novel transitions described in our work. (B) Enhanced electrical activity accompanied by loss of GABAergic interneurons inhibits the progression of intermediate progenitors towards neurogenesis and stimulate proliferation, generation of RGCs and astrogliogenesis in an environment surrounded by activated microglia. (C) Enhanced electrical activity and sustained GABAergic activity stimulate proliferation of intermediate progenitors and neuronal differentiation/survival.

Lineage progression in the adult hippocampus has been extensively studied in the last two decades using BrdU-chasing paradigms, viral-mediated, and Cre-Lox fate mapping (Aimone et al., 2014, and references therein). These studies have shown that RGCs (Type 1 cells) express GFAP, BLBP, NESTIN, SOX2, and MASH1, are slow-dividing cells, and can also be found in a quiescent state (Song, Zhong, et al., 2012). RGCs function as the founders of the neuronal lineage, generating intermediate progenitors (type 2 cells), which in turn generate type 3 cells, constrained to a future neuronal fate and expressing TBR2, DCX and NEUROD1 (Bonaguidi et al., 2011; Suh et al., 2007). Some authors also divide Type 2 cells into two subtypes - 2a and 2b, where Type 2a cells still express several markers of RGCs (BLBP, SOX2, NESTIN) but do not show radial morphologies (Steiner et al., 2006). In contrast, Type 2b express a similar set of proteins to Type 3 cells and are considered neuronal-committed progenitors. According to this gene expression pattern, we show that more than 94% of GFP+ cells are post-mitotic DCX+ neuroblasts and 6% are proliferating DCX+ cells 3 days after tamoxifen administration in the cDcx/GFP mice. These observations are also in accordance with previous data in the literature using cDcx mice (Zhang et al., 2010) or immunocytochemistry for DCX (Brown et al., 2003; Jagasia et al., 2009), and indicate that recombination is mostly restricted to Type 2b/3 cells and early neuroblasts.

Only one single GFP+ cell was observed to express the astrocyte protein GFAP 3 days after recombination. Together with our observations in the SVZ-RMS-OB system, this observation suggests that the ectopic recombination of GFAP-expressing cells is unlikely, since we failed to detect any proliferating GFP+/GFAP+ in control mice 3 days after recombination. Thus, it is reasonable to speculate that at least some of the GFAP+ cells observed in the Dcx-lineage 7 and 30 days after recombination could be generated directly or indirectly from Dcx-progenitors. These evidences may indicate that Type 2a/b progenitors are either bipotent or could regress to a more primitive stage in the lineage (Type 1/2a), when the astrogliogenic potential is recognized (Bonaguidi et al., 2011; Encinas et al., 2011). According to this latter possibility, we show that a few GFP+ cells in the SGZ maintain morphologies similar to RGL and express GFAP 30 days after recombination. The frequency of these GFP+ RGL cells is significantly increased in the DG of mice receiving KA, where we also observe an increase in astrogliogenesis. Thus, our fate-mapping experiments indicate that at least some Type 2 cells do not follow a neurogenic fate, but rather return to stages in the lineage retaining astrogliogenic potential (Figure 7A).

Neuronal activity in the hippocampus modulates progenitor cell proliferation, neuronal differentiation, and survival of newly generated granule cells (Aimone et al., 2014). We took advantage of two different models of mesial temporal lobe epilepsy (MTLE) to study how network activity could affect the cell lineage progression of DCX-expressing cells. Unilateral intrahippocampal injections of KA or PL elicit *status epilepticus* and spontaneous seizures in adult mice (Moura et al., 2019; Sperk et al., 1983; Turski et al., 1983). However, we show that these two drugs have very different effects on Dcx-lineage progression. While PL injection boosted neurogenesis in both ipsi and contralateral sides, KA injection provoked a dramatic shift in the lineage of DCX-cells towards the generation of astrocytes, especially in the ipsilateral side. Interestingly, despite the fact that neuronal activity is severely increased in the contralateral side of KA injection, DCX-cells not only normally progressed to a neuronal fate in this side but also generate a larger number of neurons.

These observations suggest that increased network activity can modulate Dcx-lineage in both directions – neurogenesis and astrogliogenesis, depending on other concomitant signals (see below). Previous work in the literature fate mapped NESTIN-expressing cells and described a similar shift in the lineage of adult hippocampal progenitors towards an astroglial fate in the ipsilateral KA-treated hippocampus (Sierra et al., 2015). We propose that Dcx-expressing Type 2b progenitors fate mapped in our study could be reversing to a NESTIN+ Type 2a and even to a Type 1/RGC state in KA-injected animals and this regress in the lineage supports the enhanced astrogliogenesis in the ipsilateral side and neurogenesis in the contralateral side (Figure 7). According to this interpretation, we observe that many cells in the Dcx-lineage show RGC-like morphologies in KA treated animals 3 days after injection.

The contradictory effects of increased network activity on Dcx-cell lineage progression in the ipsi and contralateral hippocampi of KA-injected animals could likely be explained by the different levels of GABA activity and inflammation in these regions. While the GABAergic PV+ plexus is mostly preserved in the contralateral side, the vast majority of GABAergic PV+ neurons acutely degenerate in the ipsilateral side (Bouilleret et al., 2000). GABA activity promotes RGC quiescence and enhances neuronal differentiation and survival, whereas reduced GABA activity favors progenitor self-renewal and astrogliogenesis (Dumitru et al., 2017; Song, Zhong, et al., 2012).

In contrast, increased network activity in the contralateral KA-injected side and both hippocampi of PL-injected animals could be coupled to an enhanced GABA activity, favoring neuronal differentiation and survival (Bao et al., 2017; Jagasia et al., 2009). For example, hilar BDNF infusion for two weeks in naïve animals boosts neuronal excitability and is sufficient to increase neurogenesis (Scharfman et al., 2005). In fact, GABA signaling is involved in granule cell survival and differentiation even before NMDA-glutamate pathways are activated (Jagasia et al., 2009). Yet, we cannot rule out the possibility that PL stimulates neurogenesis, at least partially, through the direct activation of muscarinic receptors (Freitas et al., 2006). Depletion of acetylcholine leads to a reduction in neurogenesis (Cooper-Kuhn et al., 2004; Kaneko et al., 2006), whereas increased cholinergic neurotransmission induced by galantamine, an acetylcholinesterase inhibitor, promotes increased neurogenesis (Kita et al., 2014). The latter effect can be partially blocked by scopolamine and the preferential M1 muscarinic receptor antagonist telenzepine (Kita et al., 2014), suggesting that progenitor cells in the hippocampus express muscarinic receptors and are affected by cholinergic activity.

Another important difference observed in the different SE models used in this study is the pattern of microglial activation/inflammation. Although increased numbers of activated microglia are present in the CA1/CA3 regions of both hippocampi of KA- and PL-injected animals, the same phenomenon was observed exclusively in the DG of KA-injected animals. These different patterns of microglial activation could help to conciliate the seemingly controversial effects of inflammation on adult neurogenesis: whereas some groups report impaired neurogenesis in the inflamed hippocampus (Ekdahl et al., 2003; Monje et al., 2003), other studies support that inflammation can contribute to the maintenance of neurogenesis (Butovsky et al., 2006). Furthermore, the observation of decreased neurogenesis only in the ipsilateral hippocampus of KA-injected animals, where increased inflammation and GABAegic neurodegeneration are also present, may suggest that inflammation *per se* is not a cause, but a consequence of neuronal and progenitor loss (Bonde et al., 2006; Sierra et al., 2015). Future experiments are required to disentangle these effects and help to establish definite causal relations between inflammation or loss of PV+ neurons with the switch in the cell lineage towards astrogliogenesis.

Collectively, our results convey a new view on the lineage progression of adult neural stem cells, indicating that intermediate progenitors can regress to more primitive states and switch the potential to generate neurons or glial cells. These alterations in cell-lineage progression correlate with microglial activation and neuronal network electric activity, with GABA potentially playing a key role in the bifurcation between neurogenesis and astrogliogenesis. Last but not least, our conflicting observations for cell lineage progression in two different models of MTLE stresses the need of more studies disentangling the precise mechanisms leading to cellular alterations in the hippocampus of epilepsy patients.

## Acknowledgments

We would like to thank Dr. Eduardo Bouth Sequerra (Brain Institute, Natal – Brazil) for providing the image showing GFP+ interneurons the olfactory bulb of cDCX/GFP animals. We also thank Laura Cocas (Santa Clara University), Adriano Tort and Draulio Araújo (Brain Institute, Natal – Brazil) for reading up and commenting on earlier versions of this manuscript, Rebecca Diniz (Laboratory of Microscopy of the Brain Institute, Brazil) for technical assistance, and Davi Drieskens for graphical design of the model in figure 7.

This work was supported by grants of the Conselho Nacional de Desenvolvimento Científico e Tecnológico (CNPq) and Fundação de Aperfeiçoamento de Pessoal de Nível Superior (CAPES) to MRC (CNPq/MS/SCTIE/DECIT # 466959/2014-1 and CAPES/PROBRAL # 23038.010291/2013-45) and to CMQ (CAPES/COFECUB grants #783/13 and #385/2019). DM was supported by a postdoctoral fellowship from CAPES/PNPD.

## Materials and Methods

Further information and requests for resources and reagents should be directed to and will be fulfilled by the Lead Contact, Marcos Costa.

## EXPERIMENTAL MODEL AND SUBJECT DETAILS

### Animals

All experiments performed involving animals were approved by the ethics committee for animals in the Federal University of Rio Grande do Norte (CEUA-UFRN) with protocol number 012/2016 conform guidelines from the regional council. For the present study, a total of 36 double-transgenic mice, with age between 8 - 12 weeks, were randomly assigned to the control or treatment (*status epilepticus* (SE) induced by kainic acid or pilocarpine) group. Mice from the lineage DCX (DCX-CRE-ER^T2^ - Stem Cell Research, Helmholtz Center Munich) were crossed to mice from the reporter lineage GFP (CAG-CAT-GFP) to generate double-transgenic DCX-CreER^T2^::CAG-CAT-GFP. The genotypes of the mutants were confirmed by PCR analysis of genomic DNA. Animals had a mixed background from C57BL/6, Swiss and Agouti. Tamoxifen (TAM) treatment of these mice restricts CreER^T2^ expression to DCX-epressing progenitors and neuroblasts in the hippocampal subgranular zone (Zhang et al., 2010) and causes random excision between multiple pairs of lox sites, leading to the expression of the reporter sequence. To achieve sparse labeling, mice were given two injections of TAM (100μg/g), in consecutive days. Sample size was calculated based in previous work considering power analysis of 80% resulting on 4 animals per group. The animals were held under standard laboratory housing conditions with a light/dark cycle of 12h each and free access to food and water. All efforts were made to minimize animal suffering.

## METHOD DETAILS

### Intrahippocampal Kainic Acid, Pilocarpine and Retroviral Injections

For unilateral intrahippocampal injections, mice were anesthetized (inhaled anesthesia mixture containing isoflurane 1.5%; 1L/min) and positioned in the stereotaxic. We injected 50nl of kainic acid (20mM in PBS) or vehicle solution (sterile PBS) using a glass micropipette attached to nanoinjector, and set to a flux of 0.5μL/min. The pipette was positioned into the right dorsal hippocampus (coordinates AP: 2.1mm; ML: 1.7; DV: 1.6mm) and left still for 5 minutes before removing to avoid liquid reflux. After KA injection, SE was characterized by clonic movements of the forelimbs, rotations or immobility. These behaviors were described before in previous studies (Riban et al., 2002) and confirmed by us through recording field potential activity in 2 animals. Animals were monitored behaviorally for seizures and the onset of SE. Following 90 minutes of SE mice were given injection of diazepam (5mg/kg, i.p.) to alleviate seizure activity. Mice were returned to their cage where they were provided free access to food and water, and monitored for weight loss and recovering in the subsequent days. Only animals that had experienced SE after KA injection, and confirmation in histological observation of granular cell dispersion were kept for further analysis.

A similar protocol was applied for Pilocarpine injection with small adaptations. Before anesthetizing the animal, methyl-scopolamine was injected (1mg/kg, i.p.) to block peripheral reactions. Coordinates and method of drug delivery were the same as KA and solution concentration and volume were 700 μg/μL and 570 μL respectively.

### Experimental Protocols

The same protocol of injections was applied for the different experimental groups with small differences in the order and timing depending on the purpose of the experiment. Each timeline is graphically presented in the results session. Tamoxifen (TAM, T-5648, Sigma Aldrich) diluted in corn oil (10mg/ml, Sigma) was administered intraperitoneally (100ug/g) twice in consecutive days a week after the SE (Vehicle group: n=3; KA group: n=5; PL group: n=3), a day before (Vehicle group: n=3; KA group: n=3;), and 5 weeks before (Vehicle group: n=4; KA group: n=4; Pilocarpine group: n=3). In the experiments with BrdU administration, it was given either by intra-peritoneal injections (50mg/kg) or diluted in the drinking water (1mg/ml).

Intrahippocampal injection of the retrovirus encoding the DsRed fluorescent reporter under control of an internal CAG promoter (pCAG-IRES-DsRed (Heinrich et al, Plos Biol, 2010) was performed following a similar protocol as for KA injection. The viral suspension (0.5 μL) was slowly injected during 30-40 min to allow for an optimal diffusion of viral particles. Viral titers used for experiments were typically in the range of 10^6^-10^9^ transducing units/mL.

### Immunohistochemistry and Microscopy

At the end of the experiments, animals were killed by intraperitoneal injection of tiopental, transcardially perfused with saline 0,9% and immediately followed by a 4% paraformadehyde solution. Brains were removed and post-fixed overnight in the same solution at 4C. They were then cryoprotected in 30% sucrose solution for 24h, and then frozen in isopentane with dry ice and stored at −80C. Brains were sectioned coronally on a cryostat at 40um and mounted to ionized slides.

Sections were incubated in the primary antibodies: rabbit anti-glial fibrillary acidic protein (GFAP; 1:1000; DAKO), chicken anti-green fluorescent protein (GFP; 1:500; Aves Labs), rabbit anti-Ctip2 (1:500; Abcam), rabbit anti-DCX (1:1000, Abcam); rat anti-BrdU (1:500, Abcam), mouse Anti-PV (1:1000, Sigma); mouse anti-NeuN (1:1000; Millipore); rabbit anti-Iba1 (1:1000, Wako). The following secondary antibodies were used: Alexa 488 goat anti-chicken, Alexa 546 goat antirabbit, Alexa 633 goat anti-rat. Immunofluorescence triple-labeling was carried out for GFP and other primary (overnight) and secondary antibodies (2 hours) incubation in permeabilization solution (PBS-TritonX 0,5%) and blocker Normal Goat Serum (NGS; 5%). To allow BrdU detection, slices were previously incubated with PFA 4% for 10 minutes and then carried with BrdU pre-treatment with sodium citrate solution 10mM at 97C for 15 min (Tang et al., 2007). After this step, immunostaining proceeded as described before.

### Quantification and statistical analysis

#### Sampling strategy

Histological analyses were performed in serial sections sampled at 1/10 (i.e., one section every 400 μm, in the anteroposterior axis). For quantitative analysis we used Stereo investigator software to count cells and to measure the areas of analyses. The subdivisions of the DG - hilar region, molecular layer (ML) and granule cell layer (GCL) were identified using the DAPI staining. For GFP quantification, a total of 4 to 5 sections were analyzed per slide and the total number of GFP cells was counted in each section. To count markers that needed co-labeling confirmation (DCX, BrdU, CTIP2, NEUN, GFAP), we captured images using a confocal laser-scanning microscope (Carl Zeiss, Jena, Germany) aiming at all GFP+ cells present in the DG. To quantify cell fates, the proportion of marker+/GFP+ or marker+/DsRed+ cells was normalized for the total number of GFP+ or DsRed+ cells, respectively. For proliferating cells and their progeny, normalization was made taking into account BrdU uptake (GFP+/BrdU+ cells).

For PV quantification, we counted PV+ positive cells spread along the hippocampus, including DG, CA3-CA2 and CA1. The slices were distributed from Bregma **−1.34 to −3.4**. All the area was identified by DAPI staining. The number of PV cells was quantified in a total of 3 to 5 sections per slide using a Zeiss Imager M.2 Apotome microscope (Carl Zeiss) with an 20x objective and the Stereoinvestigator software. For IBA1 quantification, we performed pictures of 4 areas distributed between CA1 (SR and SLM) and DG (Hilus - 2 samples). Images were acquired in 40x objective in a confocal microscope (Zeiss Examiner z.1). The number of microglial cells were distributed between 3 to 5 sections per animals in all groups (control, PL and KA). The slices were distributed from Bregma **−1.34 to −2.54.** We used ImageJ Software for analysis with MorphoLibJ integrated library and plugin. (v1.4; Legland et al., 2016; Schindelin et al., 2012). Quantification was made following methods described in Davis et al., 2017 using an adapted multistep algorithm. First, in each maximum projection of labelled microglia we applied a grey scale attribute opening filter (area minimum: 50pixels; connectivity: 8). To separate microglia soma from processes, we used an opening morphological filter (1-pixel radius octagon), before a maximum entropy threshold to segment the microglia soma from image background. We double-checked the contouring of soma and completed measuring soma area with ImageJ Analyze Particles function. Classification of microglia was made based on the morphology of cell body and processes and area measurement was used for posterior analysis. Activated microglia was defined as IBA1+ cells with enlarged cell body and processes.

Values are expressed as mean +-standard error of mean (SEM). For all comparisons one-way or two-way ANOVA (treatment group and hemisphere as factors) were performed, followed by *post-hoc* tests, if appropriate. Statistical tests were performed using GraphPad Prism version 6 for Windows GraphPad Software. Confidence interval is 95%. Differences were considered statistically significant at *p<0.05; **p<0.01; ***p<0.001.

---

**Figure S1. Experimental model for lineage tracing following intrahippocampal injection.** (A) Mosaic reconstruction of the DG of cDCX/GFP animal 30 d.a.r. showing GFP expression in granule neurons. (B) Schematic representation of the intrahippocampal administration of epileptogenic drugs in the right hippocampus. (C) Electrophysiological recording during the surgery shows self-sustained Status Epilepticus (SE) in both ipsi and contralateral hippocampi after KA injection. (D) Low power image of a coronal section of the mouse brain 30 days after KA injection. Observe the severe granule cell dispersion and CA1 degeneration in the ipsilateral side, whereas the contralateral side histology is mostly preserved. GCL - Granular Cell layer; CA1 – *Cornus Ammonis*.

**Figure S2. Cell proliferation in KA injected animals.** (A) Mean number of BrdU+ cells per section in the ipsi and contralateral hippocampi 30 days after intrahippocampal vehicle- or KA-injection. Animals were treated with BrdU in the drinking water during the first 3 days after injections (see Figure 3). (B) Quantification of total number of GFP+ cells in control, KA and PL animals. Tamoxifen was administered 7 days after intrahippocampal injections and animals were killed 38 days after recombination.

**Figure S3. Neuronal fate of GFP+ recombined cells 35 days after vehicle or KA injections.** Venn diagrams showing the number of GFP+ (green), BrdU+ (red) and CTIP2+ (blue) cells in the ipsi or contralateral hippocampi of 3 controls and 3 KA animals. Observe the small intercept between GFP and BrdU cells in all conditions. Note also the large overlap between GFP and CTIP2 in controls and in the KA contralateral side.

**Figure S4. Lack of GFP expression in striatal astrocytes following KA injection.** (A) Timeline representing experimental protocol. (B) Schematic view of the KA injection site in the striatum (red dot). (C-F) Representative slice showing the absence of GFP+ astrocytes in the striatum 35 days after tamoxifen. (C) Confirmation of injection coordinates with DAPI staining (E-F) Enlarged view of merged image. (E) Random GFP+ cells can be observed in the striatum confirming the immunohistochemistry. (F) GFAP labeling showing reactive astrocytes close to the injection site. (G-I) Coronal images of the olfactory bulb showing GFP+ interneurons in the granular and glomerular layers. No astrocytes were observed.

**Figure S5. DCX+ type 2b cells regress to earlier stages in the progenitor cell lineage**. (A) Timeline representing experimental protocol in which RGL were found. (B-D) Example of RGL-like cells in the DG of cDCX/EGFP that received tamoxifen 30 days before KA injection. Note the typical radial morphology and expression of GFAP in single plane z-projection confocal images (C,D), hallmarks of type 1 progenitors. Scale bar: 20um

**Figure S6. Retroviral-mediated tracing of intermediate progenitors confirm their gliogenic potential.** (A) Timeline presenting experimental protocol. Adult (P60-90) animals received unilateral ihpc injection of a retrovirus encoding for DsRed fluorescent reporter (pCAG-IRES-DsRed) and 4 weeks later a second ihpc injection of KA or saline (controls). Animals were perfused 4 weeks after the second injection (8 weeks after retrovirus injection). (B, C) Coronal sections of the injected DG showing DsRed+ granule neurons in saline-injected controls (B) and DsRed+ glial cells in KA-treated animals (C). The white dashed line outline the granule cell layer. (D) Quantification of the proportions of DsRed+ granule neurons and DsRed+ glial cells in saline-injected controls versus KA-treated animals.

**Figure S7. Total number of parvalbuminergic cells in whole hippocampus analysis** Quantification of the number of PV-expressing cells per mm^2^ of the whole dorsal hippocampus including DG, ML and CA regions. (n control = 4; n KA = 4; n PL = 4; Statistics: ANOVA F interaction _(2,18)_ = 3.179; p=0.0657; F side _(1,18)_ = 14.53; p=0.0013; F groups _(2,18)_ = 1.469; p = 0.2565; Tukey’s multiple comparisons test DF = 18; Adjusted p value (Ipsilateral control vs KA) = 0.0418)

## Notes

### Competing Interest Statement

The authors have declared no competing interest.

## Bibliography

Aimone, J. B., Li, Y., Lee, S. W., Clemenson, G. D., Deng, W., Gage, F. H., Jb, A., Li, Y., Sw, L., Gd, C., Deng, W., & Regulation, G. F. H. (2014). REGULATION AND FUNCTION OF ADULT NEUROGENESIS: FROM GENES TO COGNITION. Figure 1, 991–1026. https://doi.org/10.1152/physrev.00004.2014

Bao, H., Asrican, B., Li, W., Gu, B., Wen, Z., Lim, S. A., Haniff, I., Ramakrishnan, C., Deisseroth, K., Philpot, B., & Song, J. (2017). Long-Range GABAergic Inputs Regulate Neural Stem Cell Quiescence and Control Adult Hippocampal Neurogenesis. Cell Stem Cell, 21(5), 604–617.e5. https://doi.org/10.1016/j.stem.2017.10.003

Bonaguidi, M. A., Wheeler, M. A., Shapiro, J. S., Stadel, R. P., Sun, G. J., Ming, G.-L. L., & Song, H. (2011). In vivo clonal analysis reveals self-renewing and multipotent adult neural stem cell characteristics. Cell, 145(7), 1142–1155. https://doi.org/10.1016/j.cell.2011.05.024

Bonde, S., Ekdahl, C. T., & Lindvall, O. (2006). Long-term neuronal replacement in adult rat hippocampus after status epilepticus despite chronic inflammation. 23(December 2005), 965–974. https://doi.org/10.1111/j.1460-9568.2006.04635.x

Bouilleret, V., Loup, F., Kiener, T., Marescaux, C., & Fritschy, J. M. (2000). Early loss of interneurons and delayed subunit-specific changes in GABA(A)-receptor expression in a mouse model of mesial temporal lobe epilepsy. Hippocampus, 10(3), 305–324. https://doi.org/10.1002/1098-1063(2000)10:3<305::AID-HIPO11>3.0.CO;2-I

Brown, J. P., Couillard-Després, S., Cooper-Kuhn, C. M., Winkler, J., Aigner, L., & Kuhn, H. G. (2003). Transient Expression of Doublecortin during Adult Neurogenesis. Journal of Comparative Neurology. https://doi.org/10.1002/cne.10874

Butovsky, O., Ziv, Y., Schwartz, A., Landa, G., Talpalar, A. E., Pluchino, S., Martino, G., & Schwartz, M. (2006). Microglia activated by IL-4 or IFN-γ differentially induce neurogenesis and oligodendrogenesis from adult stem/progenitor cells. Molecular and Cellular Neuroscience, 31(1), 149–160. https://doi.org/10.1016/j.mcn.2005.10.006

Cooper-Kuhn, C. M., Winkler, J., & Kuhn, H. G. (2004). Decreased neurogenesis after cholinergic forebrain lesion in the adult rat. Journal of Neuroscience Research, 77(2), 155–165. https://doi.org/10.1002/jnr.20116

Danzer, S. C. (2018). Contributions of Adult-Generated Granule Cells to Hippocampal Pathology in Temporal Lobe Epilepsy: A Neuronal Bestiary. Brain Plasticity, 3(2), 169–181. https://doi.org/10.3233/bpl-170056

Davis, B. M., Salinas-Navarro, M., Cordeiro, M. F., Moons, L., & Groef, L. De. (2017). Characterizing microglia activation: A spatial statistics approach to maximize information extraction. Scientific Reports, 7(1), 1–12. https://doi.org/10.1038/s41598-017-01747-8

Deisseroth, K., Singla, S., Toda, H., Monje, M., Palmer, T. D., & Malenka, R. C. (2004). Excitation-Neurogenesis Coupling in Adult Neural Stem / Progenitor Cells. 42, 535–552. https://doi.org/10.1016/S0896-6273(04)00266-1

Dumitru, I., Neitz, A., Alfonso, J., & Monyer, H. (2017). Diazepam Binding Inhibitor Promotes Stem Cell Expansion Controlling Environment-Dependent Neurogenesis. Neuron, 94(1), 125–137.e5. https://doi.org/10.1016/j.neuron.2017.03.003

Ekdahl, C. T., Claasen, J.-H., Bonde, S., Kokaia, Z., & Lindvall, O. (2003). Inflammation is detrimental for neurogenesis in adult brain. Proceedings of the National Academy of Sciences, 100(23), 13632–13637. https://doi.org/10.1073/pnas.2234031100

Encinas, J. M., Michurina, T. V., Peunova, N., Park, J.-H. H., Tordo, J., Peterson, D. A., Fishell, G., Koulakov, A., & Enikolopov, G. (2011). Division-coupled astrocytic differentiation and age-related depletion of neural stem cells in the adult hippocampus. Cell Stem Cell, 8(5), 566–579. https://doi.org/10.1016/j.stem.2011.03.010

Encinas, J. M., & Sierra, A. (2012). Neural stem cell deforestation as the main force driving the age-related decline in adult hippocampal neurogenesis. Behavioural Brain Research, 227(2), 433–439. https://doi.org/10.1016/j.bbr.2011.10.010

Freitas, R. M., Sousa, F. C. F., Viana, G. S. B., & Fonteles, M. M. F. (2006). Effect of gabaergic, glutamatergic, antipsychotic and antidepressant drugs on pilocarpine-induced seizures and status epilepticus. Neuroscience Letters, 408(2), 79–83. https://doi.org/10.1016/j.neulet.2006.06.014

Heinrich, C., Nitta, N., Flubacher, A., Muller, M., Fahrner, A., Kirsch, M., Freiman, T., Suzuki, F., Depaulis, A., Frotscher, M., & Haas, C. A. (2006). Reelin Deficiency and Displacement of Mature Neurons, But Not Neurogenesis, Underlie the Formation of Granule Cell Dispersion in the Epileptic Hippocampus. Journal of Neuroscience, 26(17), 4701–4713. https://doi.org/10.1523/JNEUROSCI.5516-05.2006

Jagasia, R., Steib, K., Englberger, E., Herold, S., Faus-Kessler, T., Saxe, M., Gage, F. H., Song, H., & Lie, D. C. (2009). GABA-cAMP Response Element-Binding Protein Signaling Regulates Maturation and Survival of Newly Generated Neurons in the Adult Hippocampus. Journal of Neuroscience, 29(25), 7966–7977. https://doi.org/10.1523/JNEUROSCI.1054-09.2009

Kaneko, N., Okano, H., & Sawamoto, K. (2006). Role of the cholinergic system in regulating survival of newborn neurons in the adult mouse dentate gyrus and olfactory bulb. Genes to Cells, 11(10), 1145–1159. https://doi.org/10.1111/j.1365-2443.2006.01010.x

Kita, Y., Ago, Y., Higashino, K., Asada, K., Takano, E., Takuma, K., & Matsuda, T. (2014). Galantamine promotes adult hippocampal neurogenesis via M1 muscarinic and α7 nicotinic receptors in mice. The International Journal of Neuropsychopharmacology, 17(12), 1957–1968. https://doi.org/10.1017/S1461145714000613

Kralic, J. E., Ledergerber, D. A., & Fritschy, J. M. (2005). Disruption of the neurogenic potential of the dentate gyrus in a mouse model of temporal lobe epilepsy with focal seizures. European Journal of Neuroscience, 22(8), 1916–1927. https://doi.org/10.1111/j.1460-9568.2005.04386.x

Legland, D., Arganda-Carreras, I., & Andrey, P. (2016). MorphoLibJ: Integrated library and plugins for mathematical morphology with ImageJ. Bioinformatics, 32(22), 3532–3534. https://doi.org/10.1093/bioinformatics/btw413

Lévesque, M., Avoli, M., & Bernard, C. (2016). Animal models of temporal lobe epilepsy following systemic chemoconvulsant administration. Journal of Neuroscience Methods, 260, 45–52. https://doi.org/10.1016/j.jneumeth.2015.03.009

Lin, B., Coleman, J. H., Peterson, J. N., Zunitch, M. J., Jang, W., Herrick, D. B., & Schwob, J. E. (2017). Injury Induces Endogenous Reprogramming and Dedifferentiation of Neuronal Progenitors to Multipotency. Cell Stem Cell, 21(6), 761–774.e5. https://doi.org/10.1016/j.stem.2017.09.008

Lledo, P.-M., Alonso, M., & Grubb, M. S. (2006). Adult neurogenesis and functional plasticity in neuronal circuits. Nature Reviews. Neuroscience, 7(3), 179–193. https://doi.org/10.1038/nrn1867

Marín-Burgin, A., Mongiat, L. A., Pardi, M. B., Schinder, A. F., Praag, H. van, Jessberger, S., Kempermann, G., Laplagne, D. A., Lledo, P. M., Alonso, M., Grubb, M. S., Ramirez-Amaya, V., Marrone, D. F., Gage, F. H., Worley, P. F., Barnes, C. A., Clelland, C. D., Lacefield, C. O., Itskov, V Gage, F. H. (2012). Unique processing during a period of high excitation/inhibition balance in adult-born neurons. Science (New York, N.Y.), 335(6073), 1238–1242. https://doi.org/10.1126/science.1214956

Monje, M. L., Toda, H., & Palmer, T. D. (2003). Inflammatory Blockade Restores Adult Hippocampal Neurogenesis. Science, 302(5651), 1760–1765. https://doi.org/10.1126/science.1088417

Moura, D. M. S., de Sales, I. R. P., Brandão, J. A., Costa, M. R., & Queiroz, C. M. (2019). Disentangling chemical and electrical effects of status epilepticus-induced dentate gyrus abnormalities. Epilepsy and Behavior, xxxx, 106575. https://doi.org/10.1016/j.yebeh.2019.106575

Nakamura, T., Colbert, M. C., & Robbins, J. (2006). Neural crest cells retain multipotential characteristics in the developing valves and label the cardiac conduction system. Circulation Research, 98(12), 1547–1554. https://doi.org/10.1161/01.RES.0000227505.19472.69

Riban, V., Bouilleret, V., Pham-Lê, B. T., Fritschy, J. M., Marescaux, C., & Depaulis, A. (2002). Evolution of hippocampal epileptic activity during the development of hippocampal sclerosis in a mouse model of temporal lobe epilepsy. Neuroscience, 112(1), 101–111. https://doi.org/10.1016/S0306-4522(02)00064-7

Sailor, K. A., Schinder, A. F., & Lledo, P. M. (2017). Adult neurogenesis beyond the niche: its potential for driving brain plasticity. Current Opinion in Neurobiology, 42, 111–117. https://doi.org/10.1016/j.conb.2016.12.001

Scharfman, H., Goodman, J., Macleod, A., Phani, S., Antonelli, C., & Croll, S. (2005). Increased neurogenesis and the ectopic granule cells after intrahippocampal BDNF infusion in adult rats. Experimental Neurology, 192(2), 348–356. https://doi.org/10.1016/j.expneurol.2004.11.016

Schindelin, J., Arganda-Carreras, I., Frise, E., Kaynig, V., Longair, M., Pietzsch, T., Preibisch, S., Rueden, C., Saalfeld, S., Schmid, B., Tinevez, J. Y., White, D. J., Hartenstein, V., Eliceiri, K., Tomancak, P., & Cardona, A. (2012). Fiji: An open-source platform for biological-image analysis. Nature Methods, 9(7), 676–682. https://doi.org/10.1038/nmeth.2019

Sierra, A., Martín-Suárez, S., Valcárcel-Martín, R., Pascual-Brazo, J., Aelvoet, S.-A., Abiega, O., Deudero, J. J., Brewster, A. L., Bernales, I., Anderson, A. E., Baekelandt, V., Maletić-Savatić, M., & Encinas, J. M. (2015). Neuronal Hyperactivity Accelerates Depletion of Neural Stem Cells and Impairs Hippocampal Neurogenesis. Cell Stem Cell, 16(5), 488–503. https://doi.org/10.1016/j.stem.2015.04.003

Song, J., M. Christian, K., Ming, G. L., & Song, H. (2012). Modification of hippocampal circuitry by adult neurogenesis. Developmental Neurobiology, 72(7), 1032–1043. https://doi.org/10.1002/dneu.22014

Song, J., Zhong, C., Bonaguidi, M. A., Sun, G. J., Hsu, D., Gu, Y., Meletis, K., Huang, Z. J., Ge, S., Enikolopov, G., Deisseroth, K., Luscher, B., Christian, K., Ming, G., & Hongjun Song. (2012). Neuronal circuitry mechanism regulating adult quiescent neural stem cell fate decision. Nature, 489(7414), 150–154. https://doi.org/10.1038/nature11306.Neuronal

Sperk, G., Lassmann, H., Baran, H., Kish, S. J., Seitelberger, F., & Hornykiewicz, O. (1983). Kainic acid induced seizures: Neurochemical and histopathological changes. Neuroscience, 10(4), 1301–1315. https://doi.org/10.1016/0306-4522(83)90113-6

Steiner, B., Klempin, F., Wang, L., Kott, M., Kettemnmann, H., & Kempermann, G. (2006). Type-2 Cells as Link Between Glial and Neuronal Lineage in Adult Hippocampal Neurogenesis. Glia, 54, 805–814. https://doi.org/10.1002/glia

Steiner, Kronenberg, G., Jessberger, S., Brandt, M. D., Reuter, K., & Kempermann, G. (2004). Differential Regulation of Gliogenesis in the Context of Adult Hippocampal Neurogenesis in Mice. Glia, 46(1), 41–52. https://doi.org/10.1002/glia.10337

Suh, H., Consiglio, A., Ray, J., Sawai, T., D’Amour, K. A., & Gage, F. H. (2007). In Vivo Fate Analysis Reveals the Multipotent and Self-Renewal Capacities of Sox2+ Neural Stem Cells in the Adult Hippocampus. Cell Stem Cell, 1(5), 515–528. https://doi.org/10.1016/j.stem.2007.09.002

Tang, X., Falls, D. L., Li, X., Lane, T., & Luskin, M. B. (2007). Antigen-Retrieval Procedure for Bromodeoxyuridine Immunolabeling with Concurrent Labeling of Nuclear DNA and Antigens Damaged by HCl Pretreatment. Journal of Neuroscience, 27(22), 5837–5844. https://doi.org/10.1523/JNEUROSCI.5048-06.2007

Tozuka, Y., Fukuda, S., Namba, T., Seki, T., & Hisatsune, T. (2005). GABAergic Excitation Promotes Neuronal Differentiation in Adult Hippocampal Progenitor Cells. Neuron, 47, 803–815. https://doi.org/10.1016/j.neuron.2005.08.023

Turski, W. A., Cavalheiro, E. A., Schwarz, M., Czuczwar, S. J., Kleinrok, Z., & Turski, L. (1983). Limbic seizures produced by pilocarpine in rats: behavioural, electroencephalographic and neuropathological study. Behavioural Brain Research, 9(3), 315–335. http://www.ncbi.nlm.nih.gov/pubmed/6639740

van Praag, H., Christie, B. R., Sejnowski, T. J., & Gage, F. H. (1999). Running enhances neurogenesis, learning, and long-term potentiation in mice. Proceedings of the National Academy of Sciences of the United States of America, 96(23), 13427–13431. http://www.ncbi.nlm.nih.gov/pubmed/10557337

Zhang, J., Giesert, F., Kloos, K., Vogt Weisenhorn, D. M., Aigner, L., Wurst, W., & Couillard-Despres, S. (2010). A powerful transgenic tool for fate mapping and functional analysis of newly generated neurons. BMC Neuroscience, 11(1), 158. https://doi.org/10.1186/1471-2202-11-158

